# Dissecting the energetics of subunit rotation in the ribosome

**DOI:** 10.1101/514448

**Authors:** Mariana Levi, Paul C. Whitford

## Abstract

The accurate expression of proteins requires the ribosome to efficiently undergo elaborate conformational rearrangements. The most dramatic of these motions is subunit rotation, which is necessary for tRNA molecules to transition between ribosomal binding sites. While rigid-body descriptions provide a qualitative picture of the process, obtaining quantitative mechanistic insights requires one to account for the relationship between molecular flexibility and collective dynamics. Using simulated rotation events, we assess the quality of experimentally-accessible measures for describing the collective displacement of the ~ 4000-residue small subunit. For this, we ask whether each coordinate is able to identify the underlying free-energy barrier and transition state ensemble (TSE). We find that intuitive structurally-motivated coordinates (e.g. rotation angle, inter-protein distances) can distinguish between the endpoints, though they are poor indicators of barrier-crossing events, and they underestimate the free-energy barrier. In contrast, coordinates based on inter-subunit bridges can identify the TSE. We additionally verify that the committor probability for the putative TSE configurations is 0.5, a hallmark feature of any transition state. In terms of structural properties, these calculations implicate a transition state in which flexibility allows for asynchronous rearrangements of the bridges as the ribosome adopts a partially-rotated orientation. These calculations provide a theoretical foundation, upon which experimental techniques may precisely quantify the energy landscape of the ribosome.

## Introduction

By describing conformational motions in terms of diffusive movements across an energy landscape, it is possible to directly connect free-energy barriers and biomolecular kinetics.^1–3^ Within this framework, the transition state ensemble (TSE) corresponds to the set of configurations that are positioned at a saddle point on the free-energy surface (i.e. the free-energy barrier), and transition paths refer to individual barrier-crossing events.^4–8^ Since structural rearrangements are often complicated and can involve thousands of atoms, it is desirable to reduce the description to a small number of essential coordinates that preserve the dominant kinetic features. In the context of protein folding and conformational rearrangements, theoretical and experimental studies have provided techniques for identifying low-dimensional free-energy barriers.^9–20^ With these tools, along with advances in simulation techniques and computational capabilities, we are now positioned to search for appropriate one-dimensional coordinates that describe large assemblies, such as the ribosome.

The bacterial ribosome is a complex molecular assembly (2.4 MDa) comprised of three RNA molecules and over 50 proteins (Fig. 1A). During the translation of mRNA sequences into proteins, the ribosome undergoes a series of conformational rearrangements that include the relative displacement of large domains (>20,000 atoms). These motions allow tRNA and mRNA molecules to sequentially bind to three distinct ribosomal binding sites (A, P and E) on the small (30S) and large (50S) subunits.^21–28^ Each round of elongation begins with delivery of an aminoacyl-tRNA (aa-tRNA) molecule by EF-Tu. After formation of codon-anticodon interactions between the aa-tRNA and mRNA, the tRNA molecule fully binds to the ribosomal A site (i.e. accommodation) and the new amino acid is added to the nascent protein chain. Upon forming a peptide bond, the ribosome-tRNA-mRNA complex is described as being in a pre-translocation state. From this point, the tRNA molecules may spontaneously adopt hybrid-state configurations, prior to binding the next ribosomal binding site (i.e. translocation). Hybrid configurations are associated with ~ 20 − 50Å displacements of the tRNA molecules, which are facilitated by transient large-scale rotary-like motion (Fig. 1) of the ribosomal subunits.^29–33^

**Figure 1:**
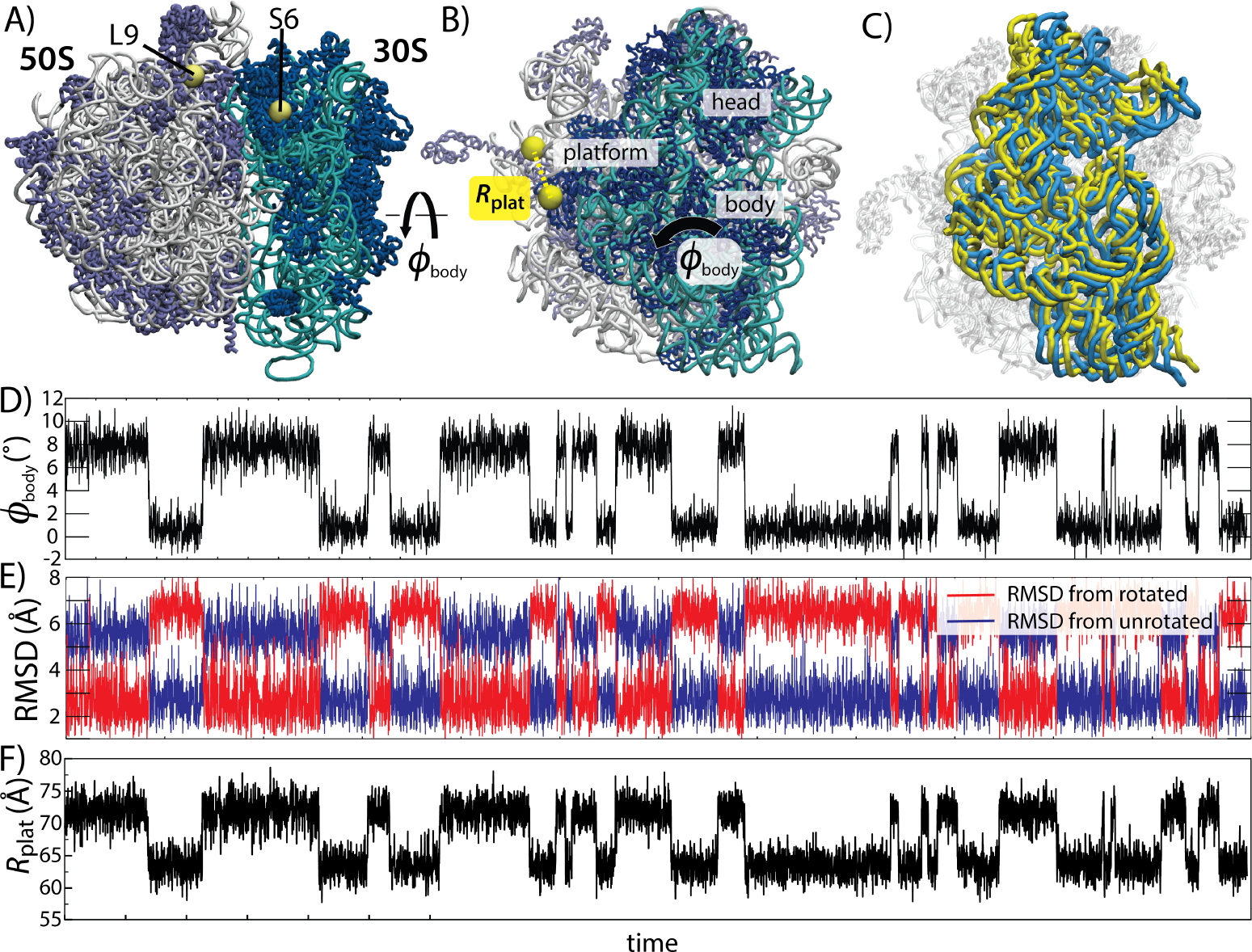
Subunit rotation in the ribosome. **A**) The 70S ribosome shown in tube representation. The 30S rRNA and proteins are shown in cyan and blue, and the 50S rRNA and proteins are colored in white and purple. During mRNA-tRNA translocation, the 30S ribosomal subunit undergoes a counterclockwise rotation, relative to the 50S subunit. This rotation may be described in terms of the Euler angle *ϕ*_body_,^40,48^ which ranges from 0 to ≈ 10°. **B)** The 30S subunit is generally described in terms of the body, platform and head regions. To monitor rotation, previous studied have placed FRET dyes on the proteins S6 (GLU41) and L9 (ASN11).^30^ Here, we will refer to the distance between these residues as *R*_plat_. **C)** Average unrotated (blue; *ϕ*_body_ ≈ 0.7°) and rotated (yellow; *ϕ*_body_ ≈ 7.8°) conformations of the small subunit rRNA, obtained from simulations. 50S subunit shown in white. All structural depictions were generated using VMD.^76^ **D-F)** Models that employ a coarse-grained representation (one bead per residue) of the ribosome can be used to simulate spontaneous interconversion events,^48^ as measured by *ϕ*_body_ (panel D), RMSD values (panel E) and *R*_plat_ (panel F).

Many studies have provided insights into the character of subunit rotation in the ribosome. Early cryo-EM reconstructions^34^ first identified a 6 − 10° counterclockwise rotation of the 30S subunit, relative to the 50S (Fig. 1A,B). Subsequent efforts refined this description by distinguishing between rotation of the 30S body and head,^35–38^ as well as tilting-like motion of the 30S head.^39,40^ In the present study, we focus specifically on 30S body rotation, which is required for rapid hybrid-formation and translocation.^41^ While the critical influence of body rearrangements is known, a physical-chemical/energetic understanding of the balance between rotation and tRNA motion is currently lacking. In order to enable experimental and theoretical investigation into this relationship, it is necessary to first identify physically-meaningful measures of rotation.

Numerous metrics have been proposed to describe subunit rearrangements in the ribosome, though most are based on static structures. For example, inertial tensor analysis of X-ray/cryo-EM structures provides a mathematically convenient coordinate system.^42^ Another strategy is to use the Euler-Rodriguez formula to characterize conformational differences in terms of single rigid-body rotations.^38^ As an extension of this, Euler angle decomposition can distinguish between rotation and tilting motions of the 30S body and head.^40^ Principal component analysis (PCA) of simulated trajectories^43–45^ and normal mode analysis^46,47^ provide measures of ribosomal flexibility, and spontaneous simulated rotation events have been isolated based on root-mean-square deviation (RMSD) calculations.^48^ In terms of experiments, single-molecule and bulk FRET assays have provided insights into the timing of subunit rotation during tRNA hybrid-formation and translocation.^29,49–51^ Even though each experimental design has been grounded in available structural data, different labeling strategies can implicate apparently contradictory dynamics.^29,49^ While simplified models have helped rationalize these differences in terms of flexibility,^48^ it is still not clear which atomic coordinates accurately report on the rate-limiting free-energy barrier. Due to this limitation, there has yet to be a study that implicates specific characteristics of the transition state ensemble associated with rotation.

In the current study, we identify structural metrics that can precisely identify the transition state ensemble and free-energy barrier of subunit rotation. For this, we analyze a recently-reported set of simulations in which hundreds of spontaneous subunit rotation events were observed.^48^ In the original study, the coarse-grained (one bead per residue) model was used to show how ribosomal flexibility can give rise to distinct relationships between individual interatomic coordinates (e.g. FRET pair distances) and rotation. Building upon this, we now adopt an energy landscape approach to assess the ability of structurally-motivated coordinates to accurately capture barrier-crossing events. We find that a collective body angle coordinate *ϕ*_body_ outperforms measures based on FRET pair distances, thought it does not identify the TSE, and it underestimates the free-energy barrier. However, we find that contact-based coordinates provide more precise descriptions of rotation events. In addition, these coordinates can reliably identify configurations for which the committor probability is 0.5, a necessary property of the TSE. From this, we are able to distinguish between bridge interactions that directly influence the kinetics, and those which primarily shift the thermodynamic balance of the system. With regards to structural characteristics, we find the TSE is associated with partial formation/breakage of specific bridges, suggests how these bridges gradually “hand off” the small subunit during rotation. Together, this analysis provides a quantitative foundation that can guide development of next-generation experimental and theoretical methods, which have the potential to uncover the precise molecular factors that govern biological assemblies. In terms of organization, we provide a more technically-oriented description under Results, while biological and experimental implications are described in the Discussion section.

## Methods

### Coarse-grained structure-based model

To explore the influence of molecular flexibility on the apparent dynamics of subunit rotation, we previously developed a coarse-grained structure-based (SMOG) model of the T. Thermophilus ribosome. The model was constructed, such that the system will undergo spontaneous rotation/back-rotation rearrangements (Fig. 1D), while accounting for molecular flexibility through an elastic network within each subunit (intrasubunit contacts). The strength of the elastic network interactions was parametrized in order for the by-residue RMSF values to be consistent with explicit-solvent simulations at 300K and with anisotropic B-factors obtained from x-ray crystallography. To allow for spontaneous large-scale transitions between the endpoint structures, intersubunit interactions were modeled using Gaussian-based Lennard-Jones-like potentials.^52^ Specifically, interface contacts that are unique to either the rotated or unrotated configuration were included as stabilizing interactions, which introduces energetic competition between the endpoints. As a result, the system spontaneously interconverts between two dominant free-energy minima that correspond to unrotated and rotated ensembles. Backbone geometry is maintained through a combination of harmonic and cosine potentials that account for bonded interactions. To prevent chain crossing, each residue was assigned an excluded volume. Non-bonded atom pairs that do not form contacts in either structure were assigned non-specific excluded volume interactions. Further details and the functional form of the coarse-grained potential can be found in the SI and Levi et al. ^48^

### MD simulations

#### Transition path analysis of equilibrium simulations

Transition path analysis was performed using a set of simulations of unrestrained rotation events. For this, molecular dynamics simulations were performed using the Gromacs (v4.5.4) software package^53,54^ with source-code extensions to allow for Gaussian-based contacts.^52^ The SMOG 2 software package (smog-server.org)^55^ was used to generate force field files. All calculations employed reduced units. Langevin Dynamics protocols were applied to ensure a constant reduced temperature of 0.67 (80, in Gromacs units). 10 independent simulations were performed, each for a minimum of 2 × 10^9^ time steps of size 0.0005. The first 2 × 10^8^ steps of each run were not analyzed to allow for equilibration of the system. For complete details, see Levi et al. ^48^

#### Committor probability calculations

For each coordinate (*R*_plat_, *ϕ*_body_ and Δ*Q*′), candidate TSE configurations were identified based on the position of the maximum in the potential of mean force (PMF, or free energy). For each of the 297 transitions paths (identified by Δ*Q*′), if the sampled frame that was closest to the TSE value was also within a narrow range of values centered about the peak (0.0705 ≤Δ*Q*′≤ 0.0715, 67.6Å ≤*R*_plat_≤ 67.61Å, or 4.19° ≤*ϕ*_body_≤ 4.195°), then it was defined as a putative TSE configuration. Following this protocol, there were 130, 125 and 110 putative TSE configurations identified for *R*_plat_, *ϕ*_body_ and Δ*Q*′. 300 independent runs were initialized from each of these configurations. Each simulation was initialized with velocities based on a Maxwell-Boltzmann distribution, and it was terminated once the system spontaneously reached one of the endpoints. Δ*Q*′ was used to determine which endpoint was reached. In total, 109,500 simulations were performed.

### Contact-based collective coordinates

To describe 30S body rotation, we specifically focused on contacts between the 30S body domain and the 50S subunit that are unique to the unrotated or rotated configurations: {*C*_unrot_}, {*C*_rot_}. The number of contacts in {*C*_unrot_} is 25 and the number of contacts in {*C*_rot_} is 68. The coordinate Δ*Q* is defined as the difference between the relative numbers of unique rotated contacts formed and unique unrotated contacts formed:

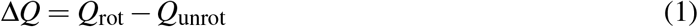

where

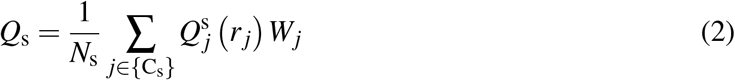

and

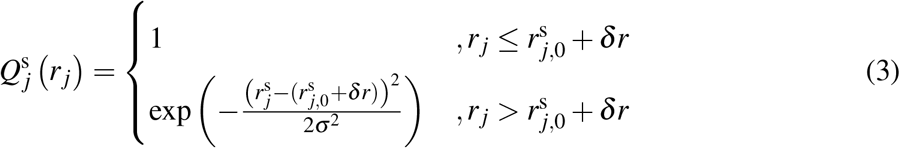

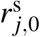is defined as the distance of contact *j* in conformation s (rotated, or unrotated), *δr* = 1Å and σ = 2Å. The endpoint configurations were defined as the average structures of the unrotated and rotated ensembles, for which *ϕ*_body_ is 0.8 and 7.8° (Fig. 1C; described in^48^). According to Equation 3, an individual contact 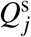 has a value of 1 if the corresponding residue pair is within 1Å of the distance found in configuration s. If the pair is extended by more than 1Å, then its contribution decays according to a Gaussian function. Equation 2 has the property that (for *W*_*j*_ = 1), if all pairs are within 1Å of their distances found in conformation s, then *Q*_s_ will reach a maximum value of 1. As all contacts specific to conformation s (rotated, or unrotated) become extended, *Q*_s_ approaches 0.

In addition to considering Δ*Q* (see SI), we also calculated Δ*Q*′(main text). The difference between these coordinates is that *W*_*j*_ is homogeneous for Δ*Q* and heterogeneous values were used for Δ*Q*′. To set the weights for Δ*Q*′, we employed a Monte-Carlo based protocol. The general strategy was to randomly select and adjust weights, and then accept the change if the maximum value of the probability of being on a transition path 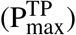 is increased. For this, the following protocol was used:

1. Initialize all weights to a uniform value: *W*_*j*_ = 1.
2. Calculate Δ*Q* for each simulated frame, as well as for the endpoint configurations: Δ*Q*_rot_, Δ*Q*_unrot_.
3. Count the number of apparent transitions between endpoints (*N*_trans_), evaluate P(TP|ΔQ) and record the maximum value 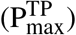.
4. Perturb the values of *W*_*j*_ (randomization details below).
5. Recalculate Δ*Q*, *N*_trans_, P(TP|ΔQ) and 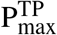.
6. The new weights are accepted if either of the following conditions are true: 1) the value of *N*_trans_ is reduced, or 2) the value of 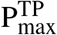 increases and *N*_trans_ does not increase.
7. Repeat steps 4-6 until 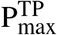 reaches a limiting value. In the current study, this cycle was repeated 30,000 times for each initialization and randomization protocol considered.

Multiple randomization schemes were considered for contact selection and weight assignments. In all optimization runs, 5000 iterations were performed where the weights of 32 randomly-selected contacts were modulated, followed by 5000 iterations in which 16, 8, 4, 2 and 1 contact weights were changed. Four different re-weighting schemes were employed: (1) toggled weights between 0 and 1, (2) randomly assign weights according to a uniform distribution between 0 and 1, (3) modulate weights by ±0.1, or (4) divided/multiple weights by 2. We found that all approaches yielded comparable values of 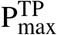. The results of protocol (1) are described in the main text. Δ*Q*′ was arbitrarily scaled by a factor of 2 after optimization. For all optimization calculations, in order to avoid overfitting during weight selection, only 1 of the 10 simulated trajectories was used in each optimization cycle. The identified coordinate was then used to analyze of the full set of simulations, which is described in the main text. This protocol was applied to each of the 10 runs, initialized at least 10 times using different random seeds.

### Functional *ϕ* value calculations

When calculating changes in free energies, which are used to determine the functional *ϕ* values (see main text), we applied free-energy perturbation techniques. For this, the original simulated data set is used to calculate the change in free energy of ensemble n according to Δ*F*_n_ = −*k*_*B*_*T* ln(⟨exp(−Δ*U*/*k*_*B*_*T*)⟩_n,0_), where ⟨⟩_n,0_ indicates the average is taken over all configurations in ensemble n for the unperturbed (i.e. “wild-type”) model and Δ*U* is the change in potential energy associated with the perturbation (i.e. “mutation”). Here, we define the perturbation as the removal of a specific intersubunit contact. We then use these perturbed free-energy values to calculate *ϕ*_1_ or *ϕ*_−1_ for each interaction. A large *ϕ* value indicates that the specific contact is formed in the TSE, whereas a low value is obtained if the contact is only formed after traversing the free-energy barrier.

## Results

To identify kinetically-relevant measures of subunit rotation, we analyzed molecular simulations in which the ribosome spontaneously interconverts between rotated and unrotated configurations. Before discussing specific results, it is important to first consider the basis of the employed model, as well as some qualitative characteristics of the simulated events. In this coarse-grained model, intrasubunit interactions are maintained through an elastic network, where the molecular flexibility is consistent with crystallographic B-factors and predictions from explicit-solvent simulations. Inter-subunit interactions are Lennard-Jones-like, allowing contacts to spontaneously form and break during rotation. By calibrating the flexibility, such that it is consistent with experimental measures, the primary utility of this model is to identify the degree to which different reaction coordinates are correlated with rotary-like rearrangements. As described elsewhere,^48^ these simple physical considerations can lead to dramatically different apparent dynamics, which previously helped rationalize apparently contradictory single-molecule measurements.

One previously-introduced measure of 30S body rotation is *ϕ*_body_,^40^ which describes the collective rearrangement of ≈400 residues in the 30S body, relative to ≈ 1400 residues in the 50S. In the current simulations, *ϕ*_body_ ranges from ~ −1° to ~ 10°, and there are abrupt transitions between ~ 1° and ~ 8° (Fig. 1D). Consistent with the dynamics along *ϕ*_body_, the values of the RMSD from the rotated and unrotated configurations are anticorrelated (Fig. 1E), and sharp transitions coincide with large changes in *ϕ*_body_. In terms of molecular structure, there is a clear distinction in domain orientation of the rotated and unrotated ensembles (Fig. 1C). Since rotation events occur spontaneously, and the model appropriately describes flexibility, we now use this data set to identify which structural measures can most closely report on the free-energy barrier associated with rotation.

### Structurally-motivated coordinates underestimate free-energy barriers

Experimentally-obtained structures can suggest intuitive metrics for describing rotation, though the apparent free-energy barrier will be influenced by the choice of reaction coordinate. To demonstrate this, we evaluated the free energy as a function of *ϕ*_body_. *ϕ*_body_ implicates a free-energy barrier of ≈ 3.4k_B_T (Fig. 2A). We next analyzed *R*_plat_, the interatomic distance between GLU41 of protein S6 and ASN11 of protein L9 (Fig. 1). These residues were considered since they have been labeled in bulk and single-molecule FRET experiments.^29–31,33,56^ Even though *R*_plat_ undergoes relatively large changes (> 10Å) during rotation, the apparent free-energy barrier is less than 2k_B_T (Fig. 2B), which is substantially smaller than the barrier obtained with *ϕ*_body_. While the dependence of the free-energy barrier height on the choice of coordinate is well documented for other systems,^8,19,57^ this apparent ambiguity for the ribosome highlights the need for systematic approaches to identify kinetically/thermodynamically-relevant coordinates for the ribosome.

**Figure 2:**
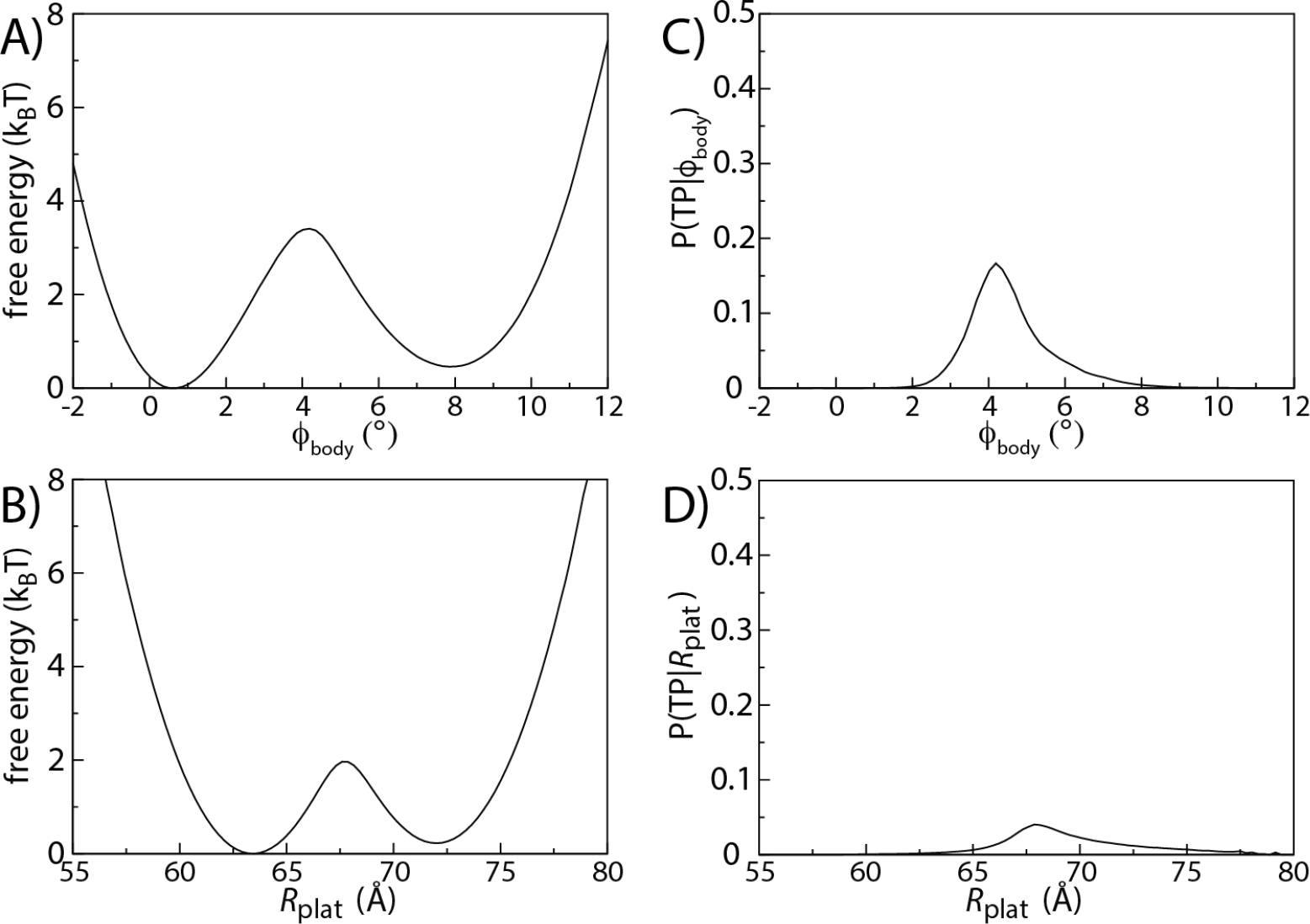
Structurally-motivated coordinates do not capture the transition state. The free energy of the ribosome as a function of **A)** *ϕ*_body_ and **B)** *R*_plat_. As expected, the height of the free-energy barrier is coordinate-dependent, where a larger barrier is obtained when using *ϕ*_body_. Consistent with the larger barrier when using *ϕ*_body_, the probability of being on a transition path *P*(TP|ρ) reaches a larger value for *ϕ*_body_ (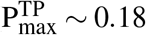, panel C) than for *R*_plat_ (panel D), indicating superior performance of *ϕ*_body_. However, 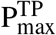 is far below the diffusion-limited value of 0.5, indicating that these coordinates do not precisely describe the transition state ensemble and free-energy barrier.

The most basic criterion to be satisfied by a good reaction coordinate is that the number of detected transitions is minimized. Reaction coordinates that understimate free-energy barriers are often associated with a large number of false positives, as seen for tRNA accommodation^58^ and hybrid-state formation.^59,60^ To evaluate the number of apparent transitions (*N*_trans_) for a given coordinate ρ, we first defined the endpoint values as the positions of the dominant free-energy minima (See Fig. 2A,B). Transition paths were then defined as segments of the trajectory in which *ρ* adopts one endpoint value and then reaches the alternate endpoint without recrossing the initial value. We find the value of *N*_trans_ depends strongly on the coordinate (Table S1). For example, *ϕ*_body_ identify 289 transitions, while *R*_plat_ appears to detect 989 events. One explanation for this high number is that *R*_plat_ may be susceptible to rapid fluctuations that manifest as false positives. To address this possibility, we calculated the survival probability *S*(τ) of the rotated and unrotated ensembles, as defined by each coordinate. If the coordinate accurately captures a two-state transition with a single rate, then *S*(τ) ~ exp(−τ/τ0), where τ_0_ = 1*/k* is the mean barrier-crossing time. We find that *S*(τ) decays according to a single exponential for *ϕ*_body_. However, *R*_plat_ yield a distribution that is more appropriately described by the sum of two exponentials (Fig. S1). Fortunately, since the decay times differ by a factor of 600-900 (Tables S2 and S3), earlier single-molecule experiments are unlikely to have detected these false event. That is, due to finite time resolution (10-100 ms), previous experiments would typically average over these rapid fluctuations.

To assess the ability of each coordinate to capture the transition state ensemble (TSE), we next analyzed the statistics of barrier-crossing events. In the diffusive regime, upon reaching any TSE, a biomolecular system will be equally likely to proceed to the “reactant” or “product” states (here, the rotated and unrotated ensembles), regardless of the most recently visited state. In other words, for any configuration **x** in the TSE, the probability of being on a transition path P(TP|**x**) will adopt a value of 0.5.^61,62^ Similarly, if a reaction coordinate *ρ* is able to precisely identify the TSE, then P(TP|ρ) will also reach a peak value of 0.5. With these considerations, we use the maximum value of P(TP|ρ), denoted 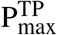, to measure the quality of each reaction coordinate. When using *ϕ*_body_, we find that 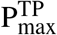 is less than 0.2 (Fig. 2C). This indicates that, even though *ϕ*_body_ can distinguish between rotated and unrotated configurations, it is a relatively poor indicator of when the system is at the TSE. Specifically, for a given value of *ϕ*_body_, the probability that the system is undergoing a transition does not exceed 20%. Similar to *ϕ*_body_, we also find that *R*_plat_ is a very weak indicator of when the system is interconverting between the unrotated and rotated ensembles (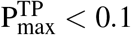; Fig. 2D). The poor and unreliable performance of these structurally-motivated coordinates further illustrates the need for a systematic approach to define effective measures of rotation.

### Subunit bridges reveal a larger free-energy barrier

Since commonly-used structurally-inspired coordinates failed to capture the transition state ensemble, we next asked whether contact-based reaction coordinates could provide a more precise description of barrier-crossing events. For this, we employed a Monte-Carlo based protocol to define collective coordinates as linear combinations of residue-residue contacts. Consistent with previous efforts,^8^ the relative weight of each intersubunit contact was iteratively adjusted. During each iteration, if the maximum probability of being on a transition path 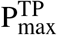 increased, then the new weights were accepted.

It should be noted that, while contact-based measures are appropriate for describing protein folding,^57,63,64^ it is not clear whether this type of metric is valuable when describing ribosome dynamics. However, previous experimental and computational results suggest that intersubunit contact (i.e. bridge interaction) dynamics will be closely related to rotation events. Structural studies have found that many intersubunit contacts transiently form and break throughout the elongation cycle,^65–68^ implying that bridge rearrangements contribute to the energetics of rotation. Explicit solvent simulations also predict that bridge energetics help determine rotation kinetics.^45^ In addition, normal mode analysis^46,47^ indicates that flexibility of the bridges predispose the ribosome to rotation. Accordingly, it is reasonable to expect that rearrangements in specific bridges may have a particularly large influence on barrier-crossing (i.e. rotation) kinetics.

To monitor the dynamics of the bridge regions, we considered differences in the number of rotated and unrotated contacts that are formed. Specifically, we calculated Δ*Q*= *Q*_rot_ − *Q*_unrot_, where *Q*_s_ describes the fraction of contacts unique to conformation s that are formed, as a function of time (Fig. 3C. See Methods for definition). To improve the precision with which one can identify the TSE, we focused our analysis on a coordinate that is defined as a linear combination of contacts, where heterogeneous weights are assigned: Δ*Q*′. To define Δ*Q*′ we applied an iterative optimization protocol in which individual weights were modified and the dynamics along each generated coordinate was analyzed. This involved randomizing the relative weight of each contact, then re-calculating the number of apparent transitions and the probability of being on a transition path, as a function of the new coordinate. At each iteration, the new weights were accepted if, 1) the number of apparent transitions decreased (i.e. fewer false positives), or 2) the number of apparent transitions was unchanged, and 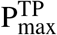 was found to increase. Using this approach, we idenetified a definition of Δ*Q*′ that yields a larger free-energy barrier and value of 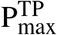. When using Δ*Q*′, we identify 297 barrier-crossing events, which is comparable to the number identified by *ϕ*_body_. Δ*Q*′ also yields a value of 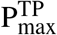 that is approximately 0.35 (Fig. 3F), which is much higher than values obtained with earlier coordinates. Finally, we find that the free-energy barrier along Δ*Q*′ is ≈ 6*k*_B_*T* (Fig. 3E), which is nearly twice as large as the barriers obtained with *ϕ*_body_ and *R*_plat_. Overall, these observations show that monitoring intersubunit contacts can more closely describe the underlying free-energy barrier and TSE than other structure-inspired coordinates.

**Figure 3:**
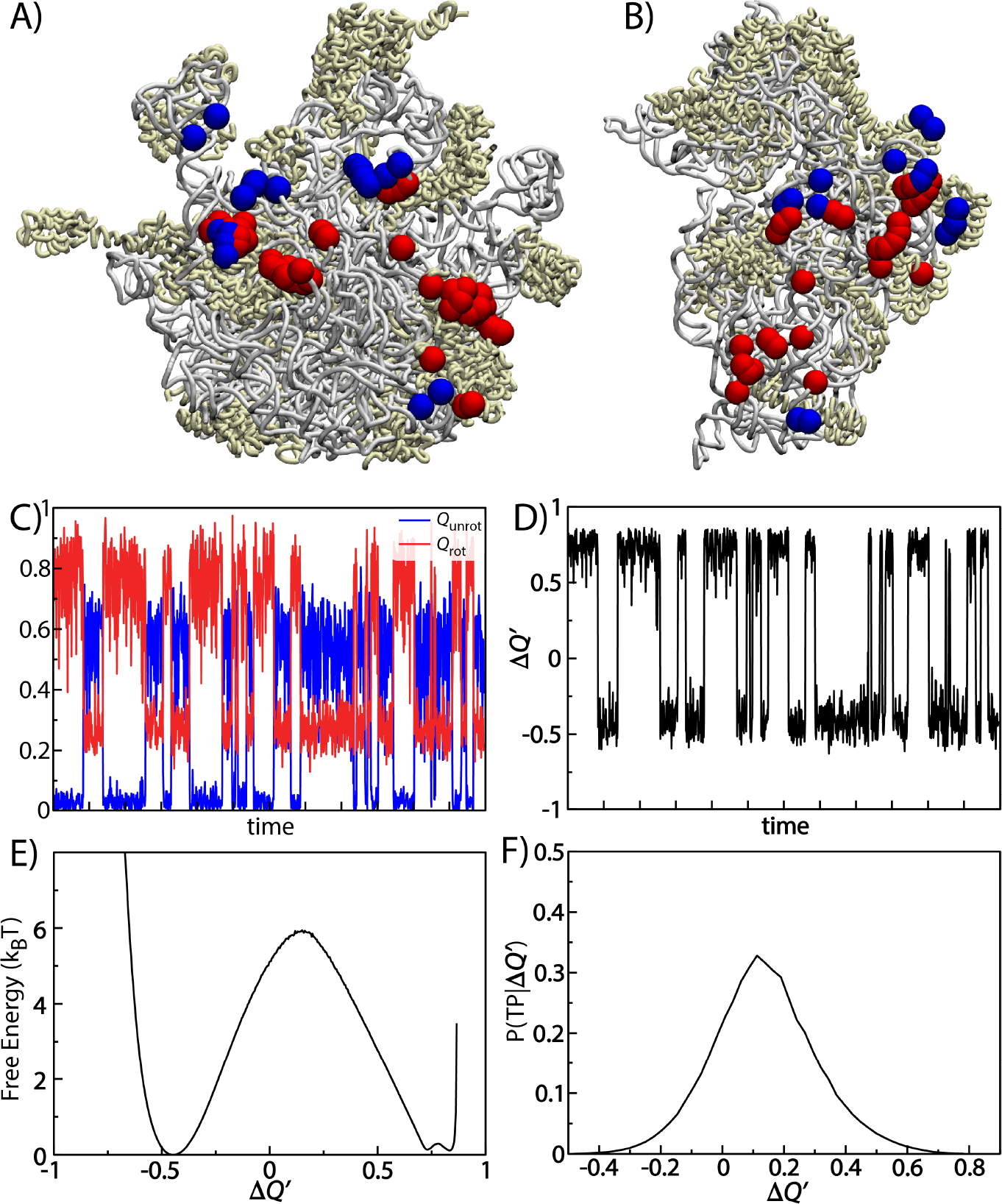
Subunit bridges implicate a larger free-energy barrier. With the relatively poor performance of structurally-motivated coordinates (Fig. 2), we considered collective reaction coordinates that monitor the subunit bridge contacts. **A)** Δ*Q* is defined as a linear combination of contacts that are unique to the rotated and unrotated configurations. Residues in the 50S that form unique rotated or unrotated contacts are shown as red and blue spheres. **B)** 30S residues that form unique rotated or unrotated contacts. To view the interface, the 30S subunit is rotated ≈ 180° about the vertical axis, relative to the 50S. **C)** Time trace of the fraction of unique unrotated contacts (*Q*_unrot_, blue) or rotated contacts (*Q*_rot_, red) for one of 10 independent simulations. **D)** Δ*Q*′, defined by a subset of bridge contacts that were selected based on a Monte-Carlo search (See Methods). **E)** The free-energy barrier obtained with Δ*Q*′ is significantly larger than other structurally-motivated coordinates (Fig. 2). **F)** The probability of being on a transition path reaches a higher value for Δ*Q*′ than *R*_plat_ and *ϕ*_body_, which indicates bridge dynamics are more precise indicators of subunit dynamics.

To further evaluate the characteristics of the TSE, we considered the committor probability of each putative TSE configuration. If the ribosome is initiated from a true transition-state configuration, then it would be equally likely to proceed to either the rotated, or unrotated free-energy minimum. In other words, the committor probability of rotating *p*_rot_ would equal 0.5. For an ideal coordinate, *p*_rot_ will be identically 0.5 for every putative TSE configuration. While that would represent a best case scenario, it is common for committor probability distributions to be narrowly centered around 0.5 for good coordinates and more broadly distributed for poor coordinates.^8^ Here, we calculated the committor probability by initializing 300 independent simulations from each putative TSE configuration (109, 500 simulations, in total). *p*_rot_ is then defined as the fraction of replicas that reached the rotated ensemble prior to the unrotated ensemble.

Committor probability calculations confirm that contact-based coordinates can isolate TSE-like configurations. That is, putative TSE configurations identified by Δ*Q*′ are most likely to have *p*_rot_ ≈ 0.5 (Fig. 4B). The distribution of *p*_rot_ values is peaked between 0.4 and 0.6, consistent with these configurations representing a TSE. This is also in agreement with the relatively high value of 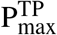 that is obtained with Δ*Q*′. In contrast, we find that *ϕ*_body_ and *R*_plat_ are unlikely to identify configurations for which *p*_rot_ ≈ 0.5 (Fig. 4A). Instead, the distributions of *p*_rot_ for these coordinates reach minimal values between 0.4-0.6, and the distributions are peaked at 0 and 1. These are very undesirable properties for any reaction coordinate, as they indicate that *ϕ*_body_ and *R*_plat_ can not reliably distinguish between the TSE and endpoint configurations.

**Figure 4:**
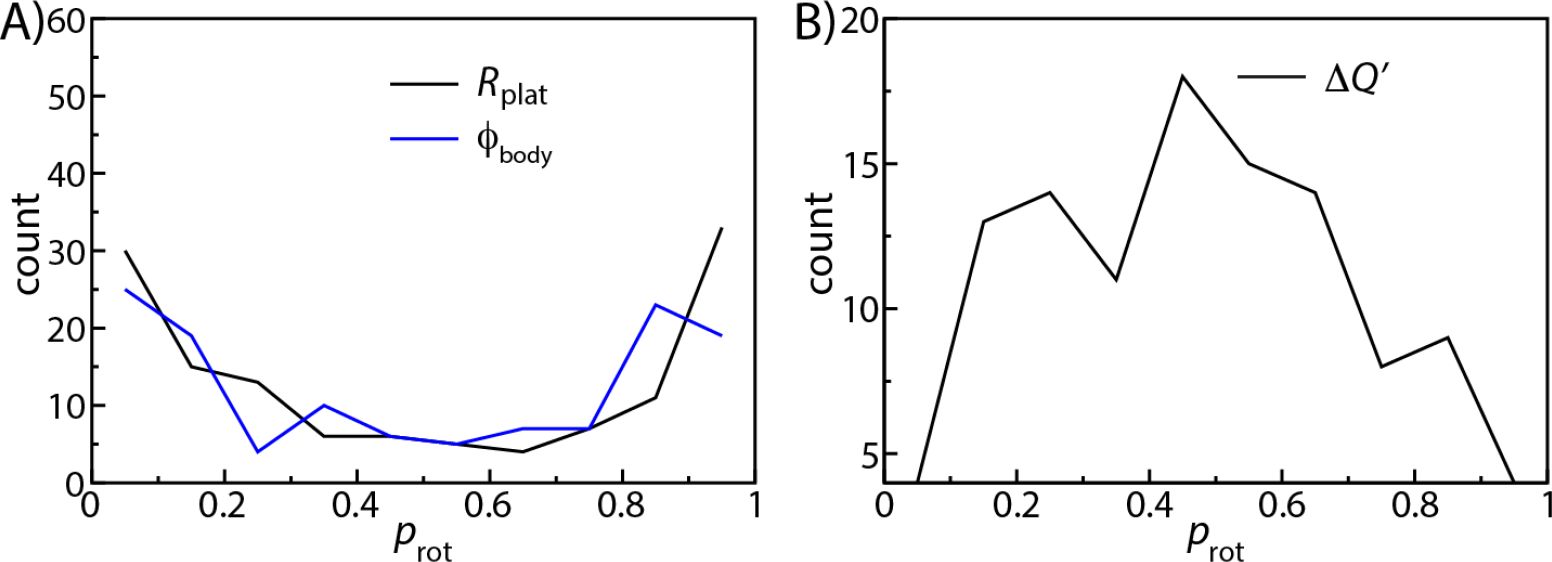
Committor probabilities of putative TSE configurations. **A**) Distribution of committor probability values, calculated for putative TSE configurations identified with *R*_plat_ and *ϕ*_body_. Very few of the identified frames have a committor probability of 0.5, indicating that the majority of the putative TSE configurations do not exhibit the characteristics of a transition state. **B)** Distribution of committor probabilities calculated for putative TSE configurations identified by Δ*Q*′. In contrast to panel A, the distribution is peaked around *p*_rot_=0.5. This indicates that Δ*Q*′ identifies TSE-like configurations, which enables the precise characterization of the underlying free-energy barrier.

Overall, the combined analysis of transition paths and committor probabilities demonstrate the superior performance of contact-based coordinates, over other structurally-motivated coordinates. The disparate performance of these two types of rotation metrics has immediate consequences on the interpretation of previous experiments, and it suggests strategies by which experiments may precisely quantify the energy landscape of subunit rotation (see Discussion).

### Functional *ϕ* values implicate asynchronous bridge rearrangements during rotation

Since Δ*Q*′ can identify configurations that are representative of the TSE, it may be used to probe the relative contribution of individual interactions to the free-energy barrier. To this end, we calculated “functional *ϕ* values,” as introduced in the study of conformational transitions of proteins.^69^ Functional *ϕ* values are an extension of traditional *ϕ*-value analysis for protein folding,^70,71^ which describes the thermodynamic effect of point mutations on the TSE. *ϕ* values have been one of the most widely used experimental measures of the energy landscapes of folding. By systematically determining changes in stabilities and kinetics, these measures have provided countless insights into the structural and energetic characteristics of folding transition states. Inspired by these successes, it is our expectation that this type of analysis strategy can prove to be equally valuable for understanding ribosome dynamics.

To apply this framework to the ribosome, we will describe subunit displacements in terms of a two-state transition between the unrotated (U) and rotated (R) ensembles:

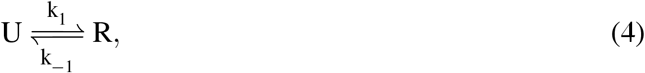

where *k*_1_ and *k*_−1_ are the rates of forward and back rotation. For interactions that stabilize forward rotation (at rate *k*_1_), the functional *ϕ* value may be defined as:

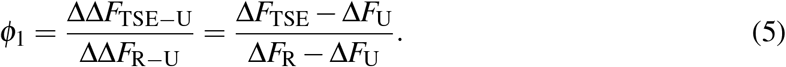

Δ*F*_TSE_, Δ*F*_U_ and Δ*F*_R_ represent the change in free energy of the TSE, the unrotated ensemble and rotated ensemble (See Fig. 5A/B) when a mutation/perturbation is introduced. According to this definition, if an interaction/residue contributes equally to the free energy of the TSE and the rotated ensemble, then *ϕ*_1_ = 1. If an interaction is only formed after the system overcomes the free-energy barrier, then *ϕ*_1_ = 0. Interactions that stabilize back-rotation can be described analogously (See Fig. S3A/B):

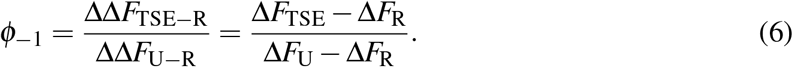

According to this construction, it is possible to distinguish between interactions that directly influence the free-energy barrier (*ϕ* ~ 1), and those which primarily stabilize the endpoints (*ϕ* ~ 0). As mentioned above, one motivating factor for adopting the *ϕ*-value framework is that it may be applied to simulations and experiments (see Discussion). Thus, this represents an avenue by which to establish a cohesive experimental/theoretical description of the ribosome’s energy landscape.

**Figure 5:**
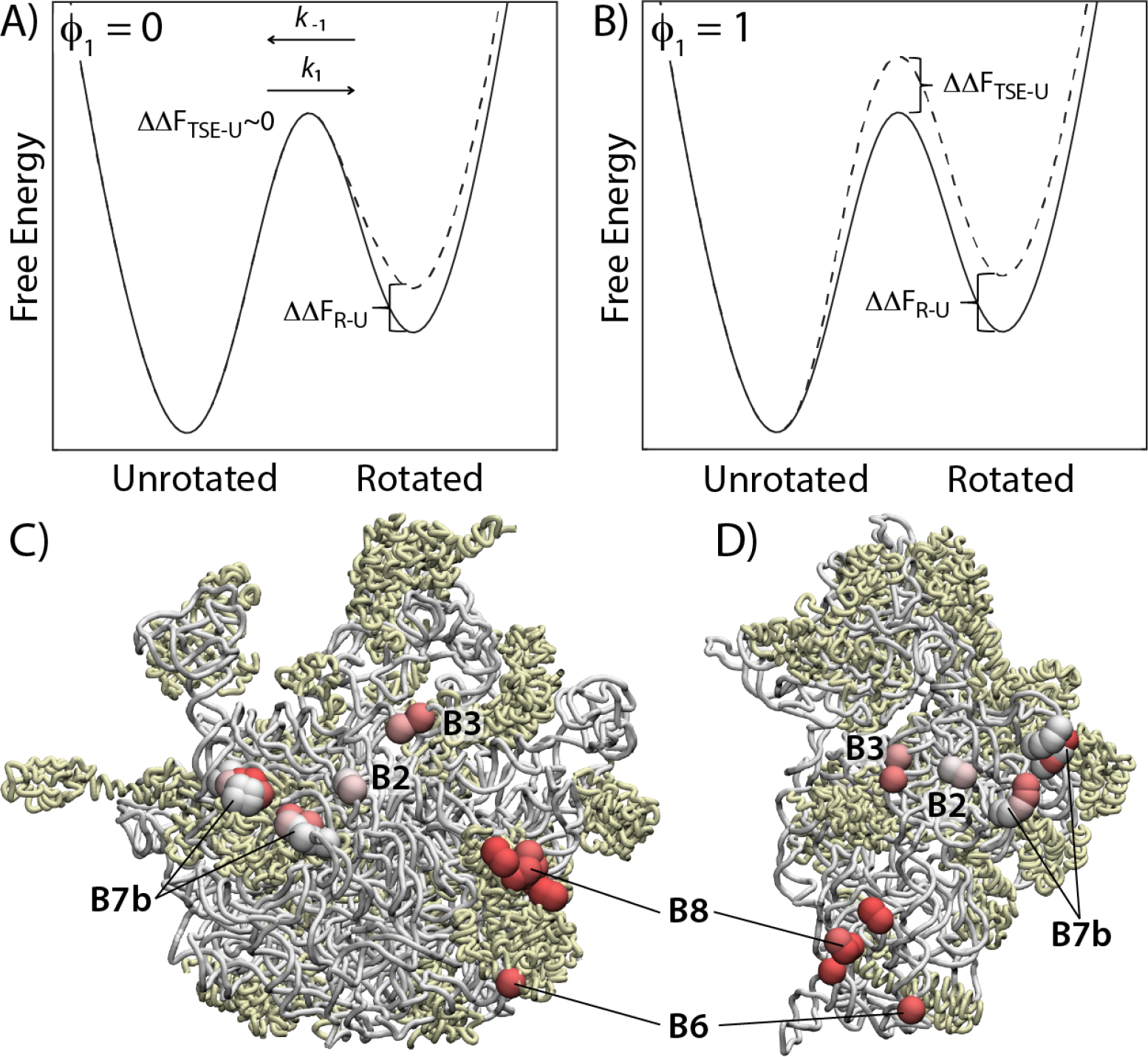
Dissecting energy landscapes through functional *ϕ* values. Introducing perturbations (e.g. mutations) to the ribosome can be used to infer characteristics of the underlying energy landscape. **A)** Schematic representation of a system where a point mutation destabilizes the rotated ensemble (ΔΔ*F*_R−U_ > 0) without impacting the transition state ensemble (ΔΔ*F*_TSE−U_ = 0). This would shift the population towards the unrotated ensemble without affecting *k*_1_. In this case, 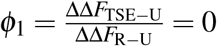. **B)** If a mutation were to destabilize the TSE and rotated ensembles to the same extent, then ΔΔ*F*_TSE−U_ and ΔΔ*F*_R−U_ would be equal, and *ϕ*_1_ would equal 1. This would reduce the forward rate and shift the population towards the unrotated ensemble. Structural representation of the 50S (panel **C**) and 30S (panel **D**), where spheres are colored by their respective value of *ϕ*_1_ (white=0, red=1). For this model, we find clusters of high *ϕ*_1_ values around residues in bridges B8 and B6, a combination of low and high values in B7b and B3, and low values for B2 residues. This illustrates how subunit bridges may progressively rearrange during the rotation process, where bridges B6 and B8 adopt their rotated orientations prior to full rotation of the subunit.

Based on our simulated trajectories, the calculated functional *ϕ* values (see Methods for details) reveal asynchronous rearrangements of the bridge regions during subunit rotation. For bridge interactions that form during the forward rotation process, there are clusters of contacts for which *ϕ*_1_ > 0.8 (Fig. 5). Residues associated with high *ϕ*_1_ values are primarily positioned within bridges B6 and B8. In particular, large values are found for contacts between protein L14 and helix 13 of the small subinit (See Table S4 for complete list of residue pairs). In contrast, contacts near/in bridge B2 have mid-range values (≈ 0.5 − 0.7) and contacts near/in bridge B7a/b tend to be associated with very low values (≈ 0 − 0.2). In terms of domain orientations, the putative transition state configurations identified by Δ*Q*′ represent a partially-rotated ensemble of configurations, where *ϕ*_body_ = 4 ± 1° (ave. ± s.d.). Together, this suggests that contacts within bridges B6 and B8 tend to form early during the rotation process, whereas B7a/b contacts tend to form as the ribosome reaches the fully-rotated ensemble. With regards to biological function, B8 interactions are known to contribute to the fidelity of tRNA selection.^72^ Our analysis suggests an extended role of B8, where it may influence fidelity while also being central to coordinating subunit rotation kinetics.

In contrast to forward rotation, we find more uniform behavior within the set of contacts that stabilize the unrotated ensemble, where almost all *ϕ*_−1_ values are less than 0.5 (Fig. S3 and Table S5). Thus, as the ribosome back-rotates, the unrotated contacts tend to form after the free-energy barrier has been crossed. While the exact value of *ϕ*_1_ and *ϕ*_−1_ are likely to depend on the details of the employed model (in simulations) and environmental/solvent conditions (in experiments), the presented calculations demonstrate how molecular flexibility can facilitate the sequential progression of bridge rearrangements during subunit rotation.

## Discussion

With continued advances in experimental and computational techniques, the field is now positioned to elucidate the detailed energetic characteristics that govern large-scale biomolecular dynamics. With decades of structural and biochemical analysis available, the ribosome is an ideal system for developing rigorous techniques for the study of molecular assemblies. To this end, we have shown how an energy landscape approach may be used to precisely describe collective dynamics. In addition to implicating general features of subunit rotation, this analysis reveals limitations in current measurements and provides design strategies for next-generation experiments, which are discussed below.

### Resolving the energy landscape with cryo-EM

Pioneering cryo-EM efforts have aimed to project large numbers of images onto low-dimensional descriptions of the configuration space, a process known as manifold embedding.^73,74^ The central idea is that one may utilize pixel-based eigenvectors to describe variations in the EM images. The eigenvectors then serve as reaction coordinates for the biological system. While this is a statistically-meaningful coordinate system, pixel covariances are likely to be more heavily influenced by larger-scale motions. Accordingly, domain orientations are likely to be clearly separated, though subunit bridge dynamics may provide only a small contribution to the final projected landscape. However, our analysis of transition events suggests that bridge rearrangements are more closely related to the free-energy barrier of rotation. As a consequence, our results indicate that orientation-dominated methods, which are sensitive to rotation angles,^37,73,74^ may underestimate the associated free-energy barriers. As these methods continue to be refined, one may envision ways in which to focus on variations in the bridge regions, which could provide a more precise measure of the underlying free-energy barrier.

### Quantifying the TSE through single-molecule measurements

As mentioned in the Results section, one motivation for calculating functional *ϕ* values is that they may be measured using currently-available single-molecule technologies. To illustrate how this may be accomplished, we will consider a residue pair in the ribosome that stabilizes the rotated conformation. For a two-state transition, the forward rate may be related to the free-energy barrier as *k*_1_ ∝ exp(−Δ*F*^†^/*k*_*B*_*T*) = exp(−(*F*_TSE_ − *F*_unrot_)/*k*_*B*_*T*). Accordingly, the forward rates for wild type 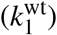 and mutant 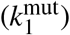 ribosomes may be used to infer the relative change in free-energies according to:

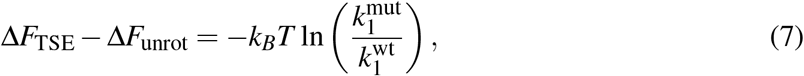

Thus, one may estimate the numerator in the definition of *ϕ*_1_ (Eq. 5) by measuring the forward rates via bulk^30–33^ or single-molecule FRET^29^ experiments. To estimate the denominator, one may also use single-molecule techniques to determine the shift in probability between the rotated and unrotated ensembles.^29^ Assuming the measured probability of occupying state n follows a Boltzmann distribution (*P*_n_ ∝ exp(−*F*_n_/*k*_B_*T*)), the denominator of Eq. 5 may be expressed as

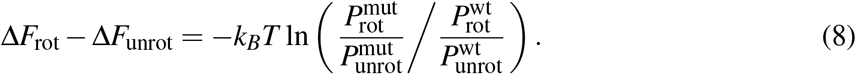

Since modern experiments can measure the relative population/occupation of each rotation state, as well as the kinetics of interconversion, functional *ϕ* values represent a means by which experimental techniques may be used to infer the energetic character of the rotation TSE. In the current study, we provide theoretical estimates of these values from a computational approach. While the exact values are likely to be model-dependent, direct comparison with experiments can allow for a cohesive and comprehensive theoretical/experimental understanding of energetics to be established.

### Deconvolving apparent dynamics and biomolecular motions

A general theme that is emerging from coordinate analysis is the need to quantitatively account for molecular flexibility when describing collective processes. This represents a significant departure from traditional rigid-body descriptions, where coordinates are implicitly assumed to be highly (albeit non-linearly) correlated. As a result of flexibility, the apparent dynamics along arbitrary coordinates can include false positives, free-energy barriers can be underestimated, and the dynamics may exhibit non-diffusive behavior. In the current study, we have found that all of these features can manifest, even for coordinates that are chosen based on knowledge of crystallographic and cryo-EM structures. In contrast, when collective coordinates are systematically analyzed, one may identify measures that are more strongly correlated with barrier crossing events.

While ideal collective coordinates are not always experimentally accessible, the presented analysis provides an avenue for inferring the scale of barriers when using non-ideal coordinates. To see how this could be accomplished, it is useful to consider two hypothetical coordinates *ρ*_1_ and *ρ*_2_, where *ρ*_1_ is an ideal coordinate that can accurately report on the barrier, and *ρ*_2_ is experimentally accessible (e.g. via single-molecule measurements). Using simulated trajectories, one may directly calculate the probability distribution along *ρ*_2_ for a given value of *ρ*_1_, *P*(*ρ*_2_|*ρ*_1_). This conditional probability distribution would represent a quantitative relationship between coordinates that accounts for the effects of molecular flexibility. An experimentally-measured distribution along *ρ*_2_ could then be related to the underlying distribution *P*(*ρ*_1_) according to 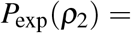 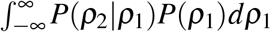. Finally, to obtain an estimate of the underlying free-energy profile, *F*(*ρ*_1_) = −*k*_*B*_*T* ln(*P*(*ρ*_1_)), one could determine *P*(*ρ*_1_) using deconvolution techniques.^75^

### Closing remarks

Calculations can provide a necessary framework that enables a wide range of experimental investigations into the energy landscape of the ribosome. Through the close integration of physical models, computation and experimental measurements, there are many opportunities available to elucidate the precise physical-chemical factors that govern large-scale dynamics in biomolecular assemblies. Inspired by studies of protein folding, we have taken the first steps towards achieving such insights for subunit rotation in the ribosome. By continuing to adopt and modify folding concepts and approaches, there is potential for future studies to elucidate the rich character of the energy landscape that controls ribosome dynamics.

## Acknowledgements

This work was supported in part by an NSF CAREER Award (Grant MCB-1350312). We acknowledge generous support provided Northeastern University Discovery Cluster and the C3DDB cluster.

## References

(1) Bryngelson, J. D.; Wolynes, P. G. Intermediates and barrier crossing in a random energy model (with applications to protein folding). The Journal of Physical Chemistry 1989, 93, 6902–6915.

(2) Bryngelson, J. D.; Onuchic, J. N.; Socci, N. D.; Wolynes, P. G. Funnels, pathways, and the energy landscape of protein folding: a synthesis. Proteins: Structure, Function, and Bioinformatics 1995, 21, 167–195.

(3) Whitford, P. C.; Onuchic, J. N. What protein folding teaches us about biological function and molecular machines. Curr. Opin. Struct. Biol. 2015, 30, 57–62.

(4) Klosek, M.; Matkowsky, B.; Schuss, Z. The Kramers problem in the turnover regime: The role of the stochastic separatrix. Berichte der Bunsengesellschaft für physikalische Chemie 1991, 95, 331–337.

(5) Du, R.; Pande, V. S.; Grosberg, A. Y.; Tanaka, T.; Shakhnovich, E. S. On the transition coordinate for protein folding. The Journal of chemical physics 1998, 108, 334–350.

(6) Bolhuis, P. G.; Chandler, D.; Dellago, C.; Geissler, P. L. Transition path sampling: Throwing ropes over rough mountain passes, in the dark. Annual review of physical chemistry 2002, 53, 291–318.

(7) Dellago, C.; Bolhuis, P.; Geissler, P. L. Transition path sampling. Advances in chemical physics 2002, 123.

(8) Best, R. B.; Hummer, G. Reaction coordinates and rates from transition paths. Proceedings of the National Academy of Sciences of the United States of America 2005, 102, 6732–6737.

(9) Das, P.; Moll, M.; Stamati, H.; Kavraki, L. E.; Clementi, C. Low-dimensional, free-energy landscapes of protein-folding reactions by nonlinear dimensionality reduction. Proceedings of the National Academy of Sciences 2006, 103, 9885–9890.

(10) Kubelka, J.; Hofrichter, J.; Eaton, W. A. The protein folding £speed limit£. Current opinion in structural biology 2004, 14, 76–88.

(11) Onuchic, J. N.; Socci, N. D.; Luthey-Schulten, Z.; Wolynes, P. G. Protein folding funnels: the nature of the transition state ensemble. Folding and Design 1996, 1, 441–450.

(12) Dobson, C. M.; Sali, A.; Karplus, M. Protein folding: a perspective from theory and experiment. Angewandte Chemie International Edition 1998, 37, 868–893.

(13) Onuchic, J. N.; Luthey-Schulten, Z.; Wolynes, P. G. Theory of protein folding: the energy landscape perspective. Annual review of physical chemistry 1997, 48, 545–600.

(14) Best, R.; Paci, E.; Hummer, G.; Dudko, O. Pulling direction as a reaction coordinate for the mechanical unfolding of single molecules. J. Phys. Chem. B 2008, 112, 5968–5976.

(15) Hyeon, C.; Thirumalai, D. Capturing the essence of folding and functions of biomolecules using coarse-grained models. Nature communications 2011, 2, 487.

(16) Chen, T.; Song, J.; Chan, H. S. Theoretical perspectives on nonnative interactions and intrinsic disorder in protein folding and binding. Current opinion in structural biology 2015, 30, 32–42.

(17) Chan, H. S.; Zhang, Z.; Wallin, S.; Liu, Z. Cooperativity, local-nonlocal coupling, and nonnative interactions: principles of protein folding from coarse-grained models. Annual review of physical chemistry 2011, 62.

(18) Grant, B. J.; Gorfe, A. A.; McCammon, J. A. Large conformational changes in proteins: signaling and other functions. Current opinion in structural biology 2010, 20, 142–147.

(19) Krivov, S. V. Is protein folding sub-diffusive? PLoS computational biology 2010, 6, e1000921.

(20) Noé, F.; Clementi, C. Collective variables for the study of long-time kinetics from molecular trajectories: theory and methods. Curr. Opin. Struct. Biol. 2017, 43, 141–147.

(21) Rodnina, M. V.; Wintermeyer, W. Fidelity of aminoacyl-tRNA selection on the ribosome: kinetic and structural mechanisms. Annual review of biochemistry 2001, 70, 415–435.

(22) Frank, J.; Gao, H.; Sengupta, J.; Gao, N.; Taylor, D. J. The process of mRNA–tRNA translocation. Proceedings of the National Academy of Sciences 2007, 104, 19671–19678.

(23) Johansson, M.; Bouakaz, E.; Lovmar, M.; Ehrenberg, M. The kinetics of ribosomal peptidyl transfer revisited. Molecular cell 2008, 30, 589–598.

(24) Wekselman, I.; Davidovich, C.; Agmon, I.; Zimmerman, E.; Rozenberg, H.; Bashan, A.; Berisio, R.; Yonath, A. Ribosome’s mode of function: myths, facts and recent results. Journal of Peptide Science 2009, 15, 122–130.

(25) Schmeing, T. M.; Ramakrishnan, V. What recent ribosome structures have revealed about the mechanism of translation. Nature 2009, 461, 1234–1242.

(26) Zaher, H. S.; Green, R. Fidelity at the molecular level: lessons from protein synthesis. Cell 2009, 136, 746–762.

(27) Korostelev, A.; Ermolenko, D. N.; Noller, H. F. Structural dynamics of the ribosome. Current opinion in chemical biology 2008, 12, 674–683.

(28) Moore, P. B.; Steitz, T. A. The roles of RNA in the synthesis of protein. Cold Spring Harbor perspectives in biology 2011, 3, a003780.

(29) Cornish, P. V.; Ermolenko, D. N.; Noller, H. F.; Ha, T. Spontaneous intersubunit rotation in single ribosomes. Mol Cell 2008, 30, 578–88.

(30) Ermolenko, D. N.; Majumdar, Z. K.; Hickerson, R. P.; Spiegel, P. C.; Clegg, R. M.; Noller, H. F. Observation of Intersubunit Movement of the Ribosome in Solution Using FRET. J. Mol. Biol. 2007, 370, 530 – 540.

(31) Ermolenko, D. N.; Spiegel, P. C.; Majumdar, Z. K.; Hickerson, R. P.; Clegg, R. M.; Noller, H. F. The antibiotic viomycin traps the ribosome in an intermediate state of translocation. Nature Structural & Molecular Biology 2007, 14, 493–7.

(32) Qin, P.; Yu, D.; Zuo, X.; Cornish, P. V. Structured mRNA induces the ribosome into a hyper-rotated state. EMBO reports 2014,

(33) Sharma, H.; Adio, S.; Senyushkina, T.; Belardinelli, R.; Peske, F.; Rodnina, M. V. Kinetics of spontaneous and EF-G-accelerated rotation of ribosomal subunits. Cell Reports 2016, 16, 2187–2196.

(34) Frank, J.; Agrawal, R. A ratchet-like inter-subunit reorganization of the ribosome during translocation. Nature 2000, 406, 318–322.

(35) Zhang, W.; Dunkle, J. A.; Cate, J. H. D. Structures of the ribosome in intermediate states of ratcheting. Science 2009, 325, 1014–7.

(36) Ratje, A. et al. Head swivel on the ribosome facilitates translocation by means of intra-subunit tRNA hybrid sites. Nature 2010, 468, 713–716.

(37) Fischer, N.; Konevega, A. L.; Wintermeyer, W.; Rodnina, M. V.; Stark, H. Ribosome dynamics and tRNA movement by time-resolved electron cryomicroscopy. Nature 2010, 466, 329–33.

(38) Mohan, S.; Donohue, J. P.; Noller, H. F. Molecular mechanics of 30S subunit head rotation. Proc. Natl. Acad. Sci. USA 2014, 111, 13325–13330.

(39) Ramrath, D. J. F.; Yamamoto, H.; Rother, K.; Wittek, D.; Pech, M.; Mielke, T.; Loerke, J.; Scheerer, P.; Ivanov, P.; Teraoka, Y.; Shpanchenko, O.; Nierhaus, K. H.; Spahn, C. M. T. The complex of tmRNA–SmpB and EF-G on translocating ribosomes. Nature 2013, 485, 526–529.

(40) Nguyen, K.; Whitford, P. C. Steric interactions lead to collective tilting motion in the ribosome during mRNA-tRNA translocation. Nat. Commun. 2016, 7, 10586–10586.

(41) Horan, L. H.; Noller, H. F. Intersubunit movement is required for ribosomal translocation. Proc. Natl. Acad. Sci. USA 2007, 104, 4881–4885.

(42) Agirrezabala, X.; Liao, H. Y.; Schreiner, E.; Fu, J.; Ortiz-Meoz, R. F.; Schulten, K.; Green, R.; Frank, J. Structural characterization of mRNA-tRNA translocation intermediates. Proceedings of the National Academy of Sciences 2012, 109, 6094–6099.

(43) Kurkcuoglu, O.; Doruker, P.; Sen, T. Z.; Kloczkowski, A.; Jernigan, R. L. The ribosome structure controls and directs mRNA entry, translocation and exit dynamics. Physical biology 2008, 5, 046005.

(44) Whitford, P. C. The ribosome’s energy landscape: Recent insights from computation. Biophysical Reviews 2015, 7, 301–310.

(45) Bock, L. V.; Blau, C.; Vaiana, A. C.; Grubmüller, H. Dynamic contact network between ribosomal subunits enables rapid large-scale rotation during spontaneous translocation. Nucleic acids research 2015, 43, 6747–6760.

(46) Tama, F.; Valle, M.; Frank, J.; Brooks, C. L. Dynamic reorganization of the functionally active ribosome explored by normal mode analysis and cryo-electron microscopy. Proc. Natl. Acad. Sci. USA 2003, 100, 9319–23.

(47) Wang, Y.; Rader, A. J.; Bahar, I.; Jernigan, R. L. Global ribosome motions revealed with elastic network model. J. Struct. Biol. 2004, 147, 302–14.

(48) Levi, M.; Nguyen, K.; Dukaye, L.; Whitford, P. C. Quantifying the Relationship between Single-Molecule Probes and Subunit Rotation in the Ribosome. Biophys. J. 2017, 113, 2777–2786.

(49) Marshall, R.; Dorywalska, M.; Puglisi, J. Irreversible chemical steps control intersubunit dynamics during translation. Proc Nat Acad Sci USA 2008, 105, 15364–9.

(50) Fei, J.; Kosuri, P.; Macdougall, D.; Gonzalez, R. Coupling of Ribosomal L1 Stalk and tRNA Dynamics during Translation Elongation. Molecular Cell 2008,

(51) Cornish, P.; Ermolenko, D.; Staple, D.; Hoang, L.; Hickerson, R.; Noller, H.; Ha, T. Following movement of the L1 stalk between three functional states in single ribosomes. Proc. Natl. Acad. Sci. USA 2009, 106, 2571–2576.

(52) Lammert, H.; Schug, A.; Onuchic, J. N. Robustness and generalization of structure-based models for protein folding and function. Proteins 2009, 77, 881–91.

(53) Lindahl, E.; Hess, B.; van der Spoel, D. GROMACS 3.0: A package for molecular simulation and trajectory analysis. J. Mol. Mod. 2001, 7, 306317.

(54) Hess, B.; Kutzner, C.; van der Spoel, D.; Lindahl, E. GROMACS 4: Algorithms for highly efficient, load-balanced, and scalable molecular simulation. J. Chem. Theory Comput. 2008, 4, 435–447.

(55) Noel, J. K.; Levi, M.; Raghunathan, M.; Lammert, H.; Hayes, R. L.; Onuchic, J. N.; Whitford, P. C. SMOG 2: A Versatile Software Package for Generating Structure-Based Models. PLoS Comput. Biol. 2016, 12, e1004794.

(56) Ermolenko, D. N.; Noller, H. F. mRNA translocation occurs during the second step of ribosomal intersubunit rotation. Nat Struct Mol Biol 2011, 18, 457–U92.

(57) Cho, S. S.; Levy, Y.; Wolynes, P. G. P versus Q: structural reaction coordinates capture protein folding on smooth landscapes. Proc. Natl. Acad. Sci. USA 2006, 103, 586–91.

(58) Noel, J. K.; Chahine, J.; Leite, V. B.; Whitford, P. C. Capturing transition paths and transition states for conformational rearrangements in the ribosome. Biophysical journal 2014, 107, 2881–2890.

(59) Nguyen, K.; Whitford, P. C. Capturing transition states for tRNA hybrid-state formation in the ribosome. The Journal of Physical Chemistry B 2016, 120, 8768–8775.

(60) Nguyen, K.; Yang, H.; Whitford, P. C. How the Ribosomal A-Site Finger Can Lead to tRNA Species-Dependent Dynamics. J. Phys. Chem. B 2017, 121, 2767–2775.

(61) Best, R. B.; Hummer, G. Reaction coordinates and rates from transition paths. Proc. Natl. Acad. Sci. USA 2005, 102, 6732–6737.

(62) Hummer, G. From transition paths to transition states and rate coefficients. The Journal of chemical physics 2004, 120, 516–523.

(63) Zhang, Z.; Chan, H. S. Transition paths, diffusive processes, and preequilibria of protein folding. Proc. Natl. Acad. Sci. USA 2012, 109, 20919–20924.

(64) Best, R. B.; Hummer, G.; Eaton, W. A. Native contacts determine protein folding mechanisms in atomistic simulations. Proc. Natl. Acad. Sci. USA 2013, 110, 17874–17879.

(65) Yusupov, M. M.; Yusupova, G. Z.; Baucom, A.; Lieberman, K.; Earnest, T. N.; Cate, J.; Noller, H. F. Crystal structure of the ribosome at 5.5 Å resolution. science 2001, 292, 883–896.

(66) Valle, M.; Zavialov, A.; Sengupta, J.; Rawat, U.; Ehrenberg, M.; Frank, J. Locking and unlocking of ribosomal motions. Cell 2003, 114, 123–134.

(67) Spahn, C. M.; Gomez-Lorenzo, M. G.; Grassucci, R. A.; Jørgensen, R.; Andersen, G. R.; Beckmann, R.; Penczek, P. A.; Ballesta, J. P.; Frank, J. Domain movements of elongation factor eEF2 and the eukaryotic 80S ribosome facilitate tRNA translocation. The EMBO journal 2004, 23, 1008–1019.

(68) Schuwirth, B. S.; Borovinskaya, M. A.; Hau, C. W.; Zhang, W.; Vila-Sanjurjo, A.; Holton, J. M.; Cate, J. H. D. Structures of the bacterial ribosome at 3.5 Å resolution. Science 2005, 310, 827–834.

(69) Whitford, P.; Miyashita, O.; Levy, Y.; Onuchic, J. Conformational transitions of adenylate kinase: Switching by cracking. J. Mol. Biol. 2007, 366, 1661–1671.

(70) Matouschek, A.; Kellis Jr, J. T.; Serrano, L.; Fersht, A. R. Mapping the transition state and pathway of protein folding by protein engineering. Nature 1989, 340, 122.

(71) Fersht, A. R.; Matouschek, A.; Serrano, L. The folding of an enzyme: I. Theory of protein engineering analysis of stability and pathway of protein folding. Journal of molecular biology 1992, 224, 771–782.

(72) Liu, Q.; Fredrick, K. Intersubunit Bridges of the Bacterial Ribosome. J. Mol. Biol. 2016, 428, 2146–2164.

(73) Dashti, A.; Schwander, P.; Langlois, R.; Fung, R.; Li, W.; Hosseinizadeh, A.; Liao, H. Y.; Pallesen, J.; Sharma, G.; Stupina, V. A.; Simon, A. E.; Dinman, J. D.; Frank, J.; Ourmazd, A. Trajectories of the ribosome as a Brownian nanomachine. Proc. Natl. Acad. Sci. USA 2014, 111, 17492–17497.

(74) Frank, J.; Ourmazd, A. Continuous changes in structure mapped by manifold embedding of single-particle data in cryo-EM. Methods 2016, 100, 61–67.

(75) Hanson, J. A.; Duderstadt, K.; Watkins, L. P.; Bhattacharyya, S.; Brokaw, J.; Chu, J.-W.; Yang, H. Illuminating the mechanistic roles of enzyme conformational dynamics. Proc Nat Acad Sci USA 2007, 104, 18055–60.

(76) Humphrey, W.; Dalke, A.; Schulten, K. VMD: Visual molecular dynamics. J. Mol. Graph. 1996, 14, 33–38.

