## Supporting Information for "Dissecting the energetics of subunit rotation in the ribosome"

### 1 Coarse-Grained Structure-based Potential

#### 1.1 Functional Form

The coarse-grained model represents each residue as a single bead of unit mass. Stabilizing interactions were defined based on rotated and unrotated configurations. Harmonic and cosine terms were introduced to maintain backbone geometry. Intrasubunit interactions (i.e. internal to 30S or 50S) were modeled according to an elastic network [1, 2, 3]. Intersubunit (between 30S and 50S) interactions were assigned Gaussian potentials [4]. To avoid chain crossing, every residue was assigned an excluded volume. In addition to the description below, a complete technical description is available elsewhere [5]. The functional form of the potential is:

$$V(\vec{x},\vec{x}^0) = \sum_{ij\in\text{bonds}} \frac{\epsilon_\text{b}}{2}\left(r_{ij}-r_{ij}^0\right)^2 + \sum_{ijk\in\text{angles}} \frac{\epsilon_\theta}{2}\left(\theta_{ijk}-\theta_{ijk}^0\right)^2 + \sum_{ijkl\in\text{dihedrals}} \epsilon_\text{d} F_\text{d}\left(\phi_{ijkl}-\phi_{ijkl}^0\right) + \sum_{ij\in\text{contacts}} U_\text{n}\left(r_{ij}^0, r_{ij}\right) + \sum_{ij\notin\text{contacts}} \epsilon_\text{nc}\left(\frac{\sigma_\text{nc}(i,j)}{r_{ij}}\right)^{12}, \tag{S1}$$

where

$$F_\text{d}(\phi) = [1-\cos(\phi)] + \frac{1}{2}[1-\cos(3\phi)]. \tag{S2}$$

$r^0_\text{ij}$, $\theta^0_\text{ijk}$ and $\phi^0_\text{ijkl}$ were assigned values based on the rotated configuration, and interresidue contacts were defined based on the Shadow algorithm [6]. Different functional forms ($U_\text{n}$) were used for distinct classes of contacts, where the energetic weight of each was $\epsilon_\text{n}$.

- $U_1$: Intrasubunit contacts - Intrasubunit contacts were included via a harmonic potential: $U_1\left(r_{ij}\right) = \frac{\epsilon_1}{2}\left(r_{ij}-r_{ij}^0\right)^2$, where the values of $r^0_\text{ij}$ are defined based on the rotated model (i.e. $r_{ij}^{0,\text{rot}}$).

- $U_2$: Interface contacts that are present in both rotated and unrotated conformations - Interface contacts that are present in both conformations were modeled via a dual-Gaussian potential [4], which has the functional form:

$$U_2\left(r_{ij}\right) = \epsilon_2\left\{\left[1+\frac{1}{\epsilon_2}\left(\frac{\sigma_\text{nc}(i,j)}{r_{ij}}\right)^{12}\right]\left[1-e^{-\frac{\left(r_{ij}-r_{ij}^{0,\text{unrot}}\right)^2}{2\sigma^2}}\right]\left[1-e^{-\frac{\left(r_{ij}-r_{ij}^{0,\text{rot}}\right)^2}{2\sigma^2}}\right]-1\right\},$$

  where $r_{ij}^{0,\text{unrot}}$ and $r_{ij}^{0,\text{rot}}$ are the pairwise distances found in the unrotated and rotated configurations. $\sigma_\text{nc}(i,j)$ is the excluded volume.

- $U_3$: Interface contacts unique to the unrotated conformation that are of a comparable distance in the rotated conformation - If a residue-residue contact is only present in the unrotated conformation, and the residue-residue distance in the rotated conformation is less than 9.8Å+1.73($r_{ij}^{0,\text{unrot}}$ − 5.9Å), then it is described by a dual-Gaussian function:

$$U_3\left(r_{ij}\right) = \epsilon_3\left\{\left[1+\frac{1}{\epsilon_3}\left(\frac{\sigma_\text{nc}(i,j)}{r_{ij}}\right)^{12}\right]\left[1-e^{-\frac{\left(r_{ij}-r_{ij}^{0,\text{unrot}}\right)^2}{2\sigma^2}}\right]\left[1-e^{-\frac{\left(r_{ij}-r_{ij}^{0,\text{rot}}\right)^2}{2\sigma^2}}\right]-1\right\}.$$

- $U_4$: Interface contacts unique to the unrotated conformation - These are defined as contacts for which the residue-residue distance in the rotated conformation is greater than 9.8Å+1.73($r_{ij}^{0,\text{unrot}}$ − 5.9Å). For these, a single-Gaussian function is assigned, which is based on the unrotated conformation:

$$U_4\left(r_{ij}\right)=\epsilon_4\left\{\left[1+\frac{1}{\epsilon_4}\left(\frac{\sigma_{\rm nc}(i,j)}{r_{ij}}\right)^{12}\right]\left[1-e^{-\frac{\left(r_{ij}-r_{ij}^{0,\rm unrot}\right)^2}{2\sigma^2}}\right]-1\right\}.$$

- $U_5$: Interface contacts that are unique to the rotated conformation - These are also described by a single-Gaussian function, which is defined by the rotated configuration:

$$U_5\left(r_{ij}\right)=\epsilon_5\left\{\left[1+\frac{1}{\epsilon_5}\left(\frac{\sigma_{\rm nc}(i,j)}{r_{ij}}\right)^{12}\right]\left[1-e^{-\frac{\left(r_{ij}-r_{ij}^{0,\rm rot}\right)^2}{2\sigma^2}}\right]-1\right\}.$$

If a residue pair is not in contact in either conformation, then an excluded volume interaction with radius $\sigma_{\rm nc}(i,j)$ was included.

#### 1.2 Specific Parameters Employed

The energetic parameters were defined as follows:

- The energetic weights were: $\epsilon=1, \epsilon_{\rm b}=100\epsilon/\text{Å}^2, \epsilon_\theta=80\epsilon/\text{rad}^2, \epsilon_{\rm d}=\epsilon, \epsilon_1=1.2\epsilon/\text{Å}^2, \epsilon_2=\epsilon, \epsilon_3=0.6\epsilon, \epsilon_4=0.27\epsilon, \epsilon_5=0.082\epsilon$.
- The elastic network spring constant $\epsilon_1$ was set to $1.2\epsilon/\text{Å}^2$. At this value, the internal subunit fluctuations (RMSF values) were comparable in the coarse-grained model and an all-atom model.
- Excluded volume distances were based on residue types: nucleic-nucleic, $\sigma_{\rm nc}(i,j)=6\text{Å}$ ; protein-protein, $\sigma_{\rm nc}(i,j)=4\text{Å}$; protein-nucleic, $\sigma_{\rm nc}(i,j)=\sqrt{6\text{Å}*4\text{Å}}=4.9\text{Å}$.
- $\sigma=1.41\text{Å}$.
- Common contacts, unique unrotated and unique rotated contacts $(U_2+U_4+U_5)$ with $\epsilon_2=\epsilon, \epsilon_4=0.26\epsilon, \epsilon_5=0.082\epsilon$.

#### 2 Supplementary Figures

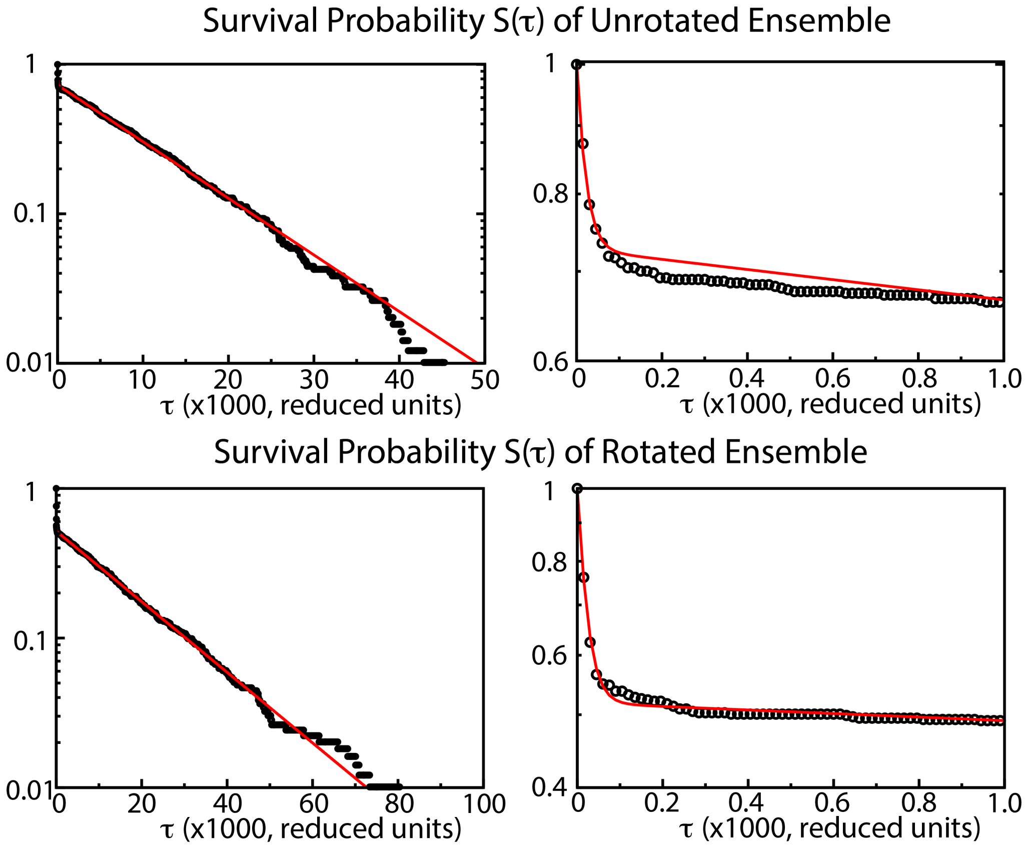

Fig. S1: Survival probabilities of the unrotated and rotated ensembles, as measured by $R_{\text{plat}}$. $S(\tau)$ is the fraction of ribosomes that remain in the unrotated (top) or rotated (bottom) ensemble after initially reaching the respective ensemble. For example, a value of 0.5 at $\tau = 100$ for state $X$ would indicate that, once the ribosome reaches state $X$, there is a 50% chance it will still be in state $X$ after 100 time units. If the coordinate describes a single barrier-crossing process (i.e. a two-state transition), then $S(\tau)$ will decay exponentially. We find $S(\tau)$ obtained with $R_{\text{plat}}$ is not well described by a single exponential fit. Rather, there is a rapid decay at low times, followed by a slow decay process. This is in contrast to the other coordinate considered in the present study, which are well described by a single exponential fit (Tables S2 and S3).

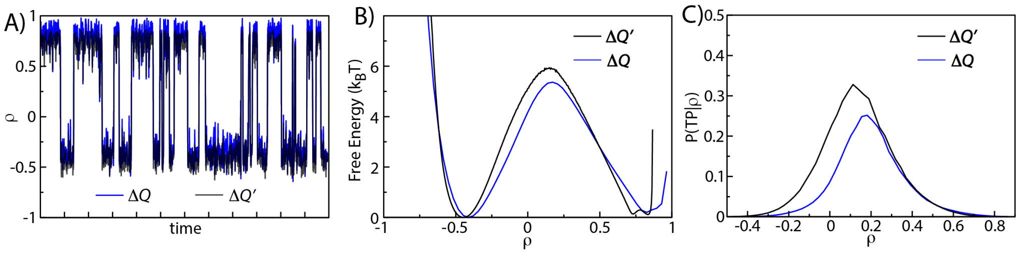

Fig. S2: **Comparison of contact-based coordinates** A) Time traces of the contact-based coordinates with homogeneous weights ($\Delta Q$) and inhomogeneous weights ($\Delta Q'$). B) Potential of mean force, calculated for the two contact-based coordinates. C) Probability of being on a transition path as a function of the contact-based coordinates.

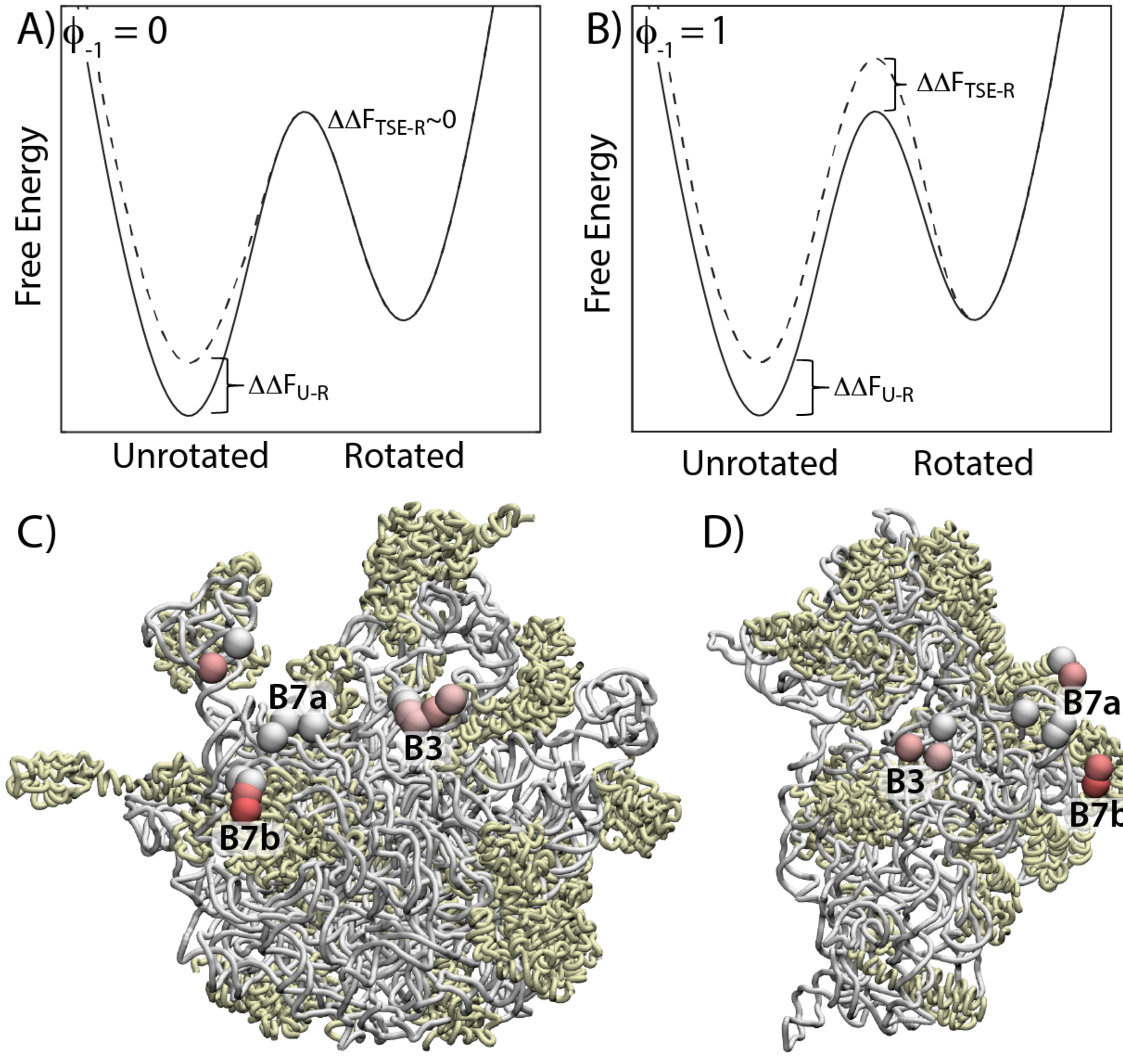

Fig. S3: **Energetic character of back-rotation.** To describe interactions that stabilize the unrotated ensemble, and thereby favor back-rotation, one may calculate $\phi_{-1}$ values. Analogous to $\phi_1$ (Fig. 5, eq. 5), $\phi_{-1}$ is defined as the ratio $\frac{\Delta\Delta F_{\mathrm{TSE-R}}}{\Delta\Delta F_{\mathrm{U-R}}}$. **A)** If a point mutation were to destabilize the unrotated ensemble, without impacting the TSE, then $\phi_{-1}$ would equal 0. This would lead to a shift in population towards the rotated ensemble without changing the rate of back-rotation. **B)** A mutation that destabilizes the unrotated ensemble and the TSE equally will reduce the rate of back-rotation, shift the population towards the rotated ensemble, and $\phi_{-1}$ will equal 1. Structure of the 50S (panel **C**) and 30S (panel **D**), where colored spheres indicate the value of $\phi_{-1}$. In contrast to $\phi_1$ (Fig. 5), most $\phi_{-1}$ values are low ($< 0.2$), though there are two residues in bridge B7b for which $\phi_{-1} > 0.7$. The generally low values of $\phi_{-1}$ indicate that most contacts specific to the unrotated ensemble only form after the ribosome has crossed the free-energy barrier.

| $\rho$ | $\rho_{\text{unrot}}$ | $\rho_{\text{rot}}$ | $N_{\text{trans}}$ | $\text{P}_{\text{max}}^{\text{TP}}$ |
|---|---|---|---|---|
| $\phi_{\text{body}}$ | 0.7° | 7.8° | 289 | 0.18 |
| $\Delta RMSD$ | -4Å | 3.1Å | 303 | 0.08 |
| $R_{\text{plat}}$ | 63.4Å | 71.9Å | 989 | 0.02 |
| $\Delta Q$ | -0.43 | 0.87 | 301 | 0.25 |
| $\Delta Q'$ | -0.24 | 0.39 | 297 | 0.32 |
| $RMSD_{\text{unrot}}, RMSD_{\text{rot}}$ | 2.5Å | 2.1Å | 335 | - |

Tab. S1: Number of apparent transitions ($N_{\text{trans}}$) and maximum probability to be on a transition path ($\text{P}_{\text{max}}^{\text{TP}}$) for each coordinate $\rho$. Endpoint values for each coordinate were defined as the positions of the minima in the free energy, or potential of mean force (PMF). All coordinates yielded a comparable number of events, except for $R_{\text{plat}}$, which overestimated the number of transitions.

| $\rho$ | $a_0$ | $\tau_0$ | $a_1$ | $\tau_1$ |
|---|---|---|---|---|
| $\phi_{\text{body}}$ | 149.993 | 25938.9 | - | - |
| $\Delta RMSD$ | 156.164 | 24950.5 | - | - |
| $R_{\text{plat}}$ | 136.976 | 20.4159 | 359.264 | 11460.1 |
| $\Delta Q$ | 154.653 | 25209.9 | - | - |
| $\Delta Q'$ | 151.273 | 25547.9 | - | - |

Tab. S2: Exponential fits to the unnormalized survival probability $N_{\text{trans}} * S(\tau)$ for the unrotated ensemble (i.e. forward rotation). Each was fit to either a single exponential $a_0 \exp(-t/\tau_1)$ or a double exponentials $a_0 \exp(-\tau/\tau_0) + a_1 \exp(-\tau/\tau_1)$. The correlation coefficient for all fits was greater than 0.99. The decay time of the probability was consistent for all coordinates, except for $R_{\text{plat}}$, which exhibited biphasic behavior.

| $\rho$ | $a_0$ | $\tau_0$ | $a_1$ | $\tau_1$ |
|---|---|---|---|---|
| $\phi_{\text{body}}$ | 145.4 | 32621.7 | - | - |
| $\Delta RMSD$ | 145.7 | 32577.6 | - | - |
| $R_{\text{plat}}$ | 238.3 | 21.3 | 256.7 | 18402.9 |
| $\Delta Q$ | 145.9 | 32549.3 | - | - |
| $\Delta Q'$ | 145.6 | 32602.2 | - | - |

Tab. S3: Exponential fits to the unnormalized survival probability $N_{\text{trans}} * S(\tau)$ for the rotated ensemble (i.e. back-rotation). Each was fit to either a single exponential $a_0 \exp(-t/\tau_1)$ or a double exponentials $a_0 \exp(-\tau/\tau_0) + a_1 \exp(-\tau/\tau_1)$. The correlation coefficient for all fits was greater than 0.99. The decay time of the probability was consistent for all coordinates, except for $R_{\text{plat}}$, which exhibited biphasic behavior.

| 50S residue | 30S body residue | bridge | $\phi_1$ |
|---|---|---|---|
| A1939(H69) | G1468(h44) | B2b | 0.552649 |
| U1862(H68) | C1489(h44) | B2b | 0.407269 |
| U1863(H68) | G1490(h44) | B2b | 0.317262 |
| ALA1(L2) | G753(h23) | B2c | 0.058399 |
| ALA1(L2) | U752(h24) | B2c | 0.114611 |
| ALA198(L2) | G753(h23) | B2c | 0.431472 |
| ASN202(L2) | G754(h23) | B2c | 0.670872 |
| ASP199(L2) | G754(h23) | B2c | 0.201140 |
| U1849(H68) | G755(h22) | B2c | 0.711423 |
| A1932(H71) | C1388(h27) | B3 | 0.697377 |
| ALA126(L19) | G1422(h44) | B6 | 0.811826 |
| ARG39(L19) | G150(h8) | B6/B8 | 0.860965 |
| ARG97(L14) | C329(h13) | B6/B8 | 0.862288 |
| PRO136(L2) | A692(h23) | B7a | 0.016734 |
| ARG167(L2) | C661(h23) | B7a/B7b | 0.000371 |
| GLU168(L2) | G662(h22) | B7a/B7b | 0.194957 |
| ALA198(L2) | G754(h23) | B7b | 0.233614 |
| ARG167(L2) | G690(h23) | B7b | 0.434255 |
| ASP199(L2) | G753(h23) | B7b | 0.210080 |
| GLN163(L2) | A692(h23) | B7b | 0.774223 |
| GLN165(L2) | C659(h22) | B7b | 0.020550 |
| GLN165(L2) | C660(h22) | B7b | 0.003080 |
| GLN165(L2) | C661(h23) | B7b | 0.030066 |
| GLN165(L2) | G690(h23) | B7b | 0.868710 |
| GLU168(L2) | C661(h23) | B7b | 0.231252 |
| GLY138(L2) | A692(h23) | B7b | 0.101925 |
| GLY138(L2) | G691(h23) | B7b | 0.140591 |
| GLY166(L2) | C660(h23) | B7b | 0.003167 |
| GLY166(L2) | C661(h23) | B7b | 0.000739 |
| GLY166(L2) | G690(h23) | B7b | 0.833407 |
| GLY169(L2) | G662(h23) | B7b | 0.007035 |
| ILE173(L2) | C660(h23) | B7b | 0.051996 |
| ILE173(L2) | C661(h22) | B7b | 0.051789 |
| LYS4(L2) | G751(h23) | B7b | 0.216422 |
| LYS4(L2) | U752(h24) | B7b | 0.171534 |
| PRO136(L2) | G691(h23) | B7b | 0.002404 |
| THR139(L2) | A692(h23) | B7b | 0.017587 |
| THR139(L2) | G693(h23) | B7b | 0.070954 |
| VAL137(L2) | A692(h23) | B7b | 0.270831 |
| VAL137(L2) | C660(h23) | B7b | 0.000137 |
| VAL137(L2) | G690(h23) | B7b | 0.055503 |
| VAL137(L2) | G691(h23) | B7b | 0.072633 |
| VAL2(L2) | G753(h23) | B7b | 0.108264 |
| VAL2(L2) | U752(h24) | B7b | 0.129072 |
| ALA118(L14) | G339(h13) | B8 | 0.867381 |
| ARG107(L14) | G338(h13) | B8 | 0.843393 |
| ASN13(L14) | A330(h13) | B8 | 0.890954 |
| GLU36(L19) | G338(h13) | B8 | 0.885432 |
| LYS113(L14) | C337(h13) | B8 | 0.757887 |
| MET112(L14) | C337(h13) | B8 | 0.739494 |
| MET112(L14) | G338(h13) | B8 | 0.800218 |
| SER116(L14) | G338(h13) | B8 | 0.819964 |
| SER116(L14) | G339(h13) | B8 | 0.752391 |
| VAL115(L14) | G338(h13) | B8 | 0.851748 |

Tab. S4: Functional $\phi$ values for contacts that stabilize the rotated conformation.

| **50S residue** | **30S body residue** | **bridge** | $\phi_{-1}$ |
|---|---|---|---|
| C2166(H78) | ALA65(S11) | - | 0.059904 |
| G1941(H71) | G771(h24) | - | 0.261159 |
| G1942(H71) | G771(h24) | - | 0.215936 |
| A1932(H71) | U1469(h44) | B3 | 0.526516 |
| A1933(H71) | U1469(h44) | B3 | 0.459263 |
| A1939(H71) | G1491(h44) | B3 | 0.461192 |
| C1940(H71) | G1491(h44) | B3 | 0.433246 |
| A1876(H68) | C661(h23) | B7a | 0.005529 |
| A1876(H68) | G662(h23) | B7a | 0.011369 |
| A1877(H68) | A682(h23) | B7a | 0.222112 |
| ARG167(L2) | ASP83(S6) | B7a | 0.014711 |
| C1914(H69) | A682(h23) | B7a | 0.115160 |
| C1915(H69) | A682(h23) | B7a | 0.150527 |
| G1878(H68) | A682(h23) | B7a | 0.086314 |
| ILE124(L2) | ILE81(S6) | B7a | 0.817356 |
| PRO136(L2) | ASP83(S6) | B7a/B7b | 0.687270 |
| PRO136(L2) | ILE81(S6) | B7a/B7b | 0.184179 |
| VAL137(L2) | ASP83(S6) | B7a/B7b | 0.169624 |

Tab. S5: Functional $\phi$ values for interface contacts that stabilize the unrotated conformation.

#### References

[1] Tirion, M. Large amplitude elastic motions in proteins from a single-parameter, atomic analysis. *Phys. Rev. Lett.* **77**, 1905–1908, 1996.

[2] Tama, F.; Valle, M.; Frank, J.; Brooks, C. L. Dynamic reorganization of the functionally active ribosome explored by normal mode analysis and cryo-electron microscopy. *Proc. Natl. Acad. Sci. USA* **100**, 9319–23, 2003.

[3] Wang, Y.; Rader, A. J.; Bahar, I.; Jernigan, R. L. Global ribosome motions revealed with elastic network model. *J. Struct. Biol.* **147**, 302–314, 2004.

[4] Lammert, H.; Schug, A.; Onuchic, J. N. Robustness and generalization of structure-based models for protein folding and function. *Prot. Struct. Func. Bioinfo.* **77**, 881–891, 2009.

[5] Levi, M.; Nguyen, K.; Dukaye, L.; Whitford, P. C. Quantifying the relationship between single-molecule probes and subunit rotation in the ribosome. *Biophys. J.* **113**, 2777–2786, 2017.

[6] Noel, J. K.; Whitford, P. C.; Onuchic, J. N. The shadow map: A general contact definition for capturing the dynamics of biomolecular folding and function. *J. Phys. Chem. B* **116**, 8692–8702, 2012.
